# Transgenerational plasticity reflects local adaptation, but is not always advantageous for fitness

**DOI:** 10.1101/2025.11.23.690023

**Authors:** Haley A. Branch, Dylan R. Moxley, Amy L. Angert

## Abstract

Transgenerational plasticity (TGP) could contribute to phenotypic differences as much as allelic variation and within-generation plasticity. Here we use northern and southern populations of *Mimulus cardinalis* collected prior to and following a severe drought and grew plants in a multi- factorial experiment across three generations to tease apart allelic, TGP, parental effects, and plasticity. Where grand-offspring are likely to experience similar stressors as grandparents, TGP is often considered advantageous; our results show no such advantage. Instead, we find a TGP pattern analogous to local adaptation. Regardless of current environment, grand-offspring exhibit increased fitness when grandparents were exposed to environments that matched historical conditions and reduced fitness when exposed to foreign environments. We also find the evolutionary loss of TGP; prior to drought, northern flowering time and southern pollen viability were significantly affected by grandparent treatment but following four years of drought, grandparent treatment no longer affected these reproductive traits. Lastly, we find that TGP can have a greater effect on fitness than allelic change, parental environment, and within-generation plasticity. Our results indicate an additional way that phenotypes arise, sheds light on the potential for how rapid evolution might occur, and has significant implications for how we design evolutionary ecology experiments.

## Introduction

Transgenerational plasticity (TGP) could be a critical factor in eliciting stress responses and rapid adaptation.^1^ In contrast to within-generation plasticity that evolves in variable climates, TGP could be advantageous when climates are predictable across generations.^2–5^ Studies suggest that TGP could help buffer populations from climate change, giving descendants an advantage when experiencing the same stress as their ancestors.^6^ TGP in the form of epigenetic modifications have been shown to increase gene mutation rate and the capacity to increase the rate of evolution.^7^ Evolution may even act on allelic variation that causes TGP instead of allelic variation in the loci underlying a trait.^7^ Alternatively, TGP could be maladaptive with rapid change, if phenotypes expressed are responding to past climates rather than present ones,^4,8^ or if stress accumulation occurs.^9,10^ However, the role that non-genetic inheritance plays in rapid evolutionary responses is not well understood, particularly in wild populations of plants, yet it will likely be increasingly more important in the future as climate change intensifies.

Here we define TGP as non-genetic inheritance by which environmentally induced phenotypic change is inherited across *more than two* generations (i.e., grandparent to grand-offspring). Most studies on non-genetic inheritance have focused on parental effects across *two generations* (parent to offspring phenotypes, parental effects).^5^ A recent review of epigenetics studies (across plants and animals) found only 11% examined more than two generations.^11^ Furthermore, studies that go beyond two generations have focused primarily at the molecular level, often using inbred lines or clonal organisms for tractability.^5,12^ Therefore, empirical evidence about the relative importance of TGP compared to within-generation plasticity, parental effects, and allelic variation for determining phenotypic diversity in wild populations is limited.^5^

Here we conducted a multi-factorial stress (wet and dry) experiment with *Mimulus cardinalis* (scarlet monkeyflower) to investigate population-level differences in the lasting phenotypic consequences of environmental stress across *three* generations (Fig. 1B). Using the resurrection approach where seed populations from ancestors and descendants are refreshed and compared in a common garden,^13^ we also assess how TGP interacts with rapid evolution during a severe drought (Fig. 1A). We found previously that populations from climatically distinct regions evolved rapidly in response to this climatic event, largely towards dehydration avoidance.^14,15^ However, we also observed lagging phenotypic change from climates from two-years prior, particularly in the north, suggestive of a grandparent-derived effect.^8^ To investigate whether this lagging phenotypic effect is caused by TGP, we examine specifically whether the grandparental environment affects grand-offspring (referred hereafter as offspring) phenotypes and responses.

**Figure 1.**
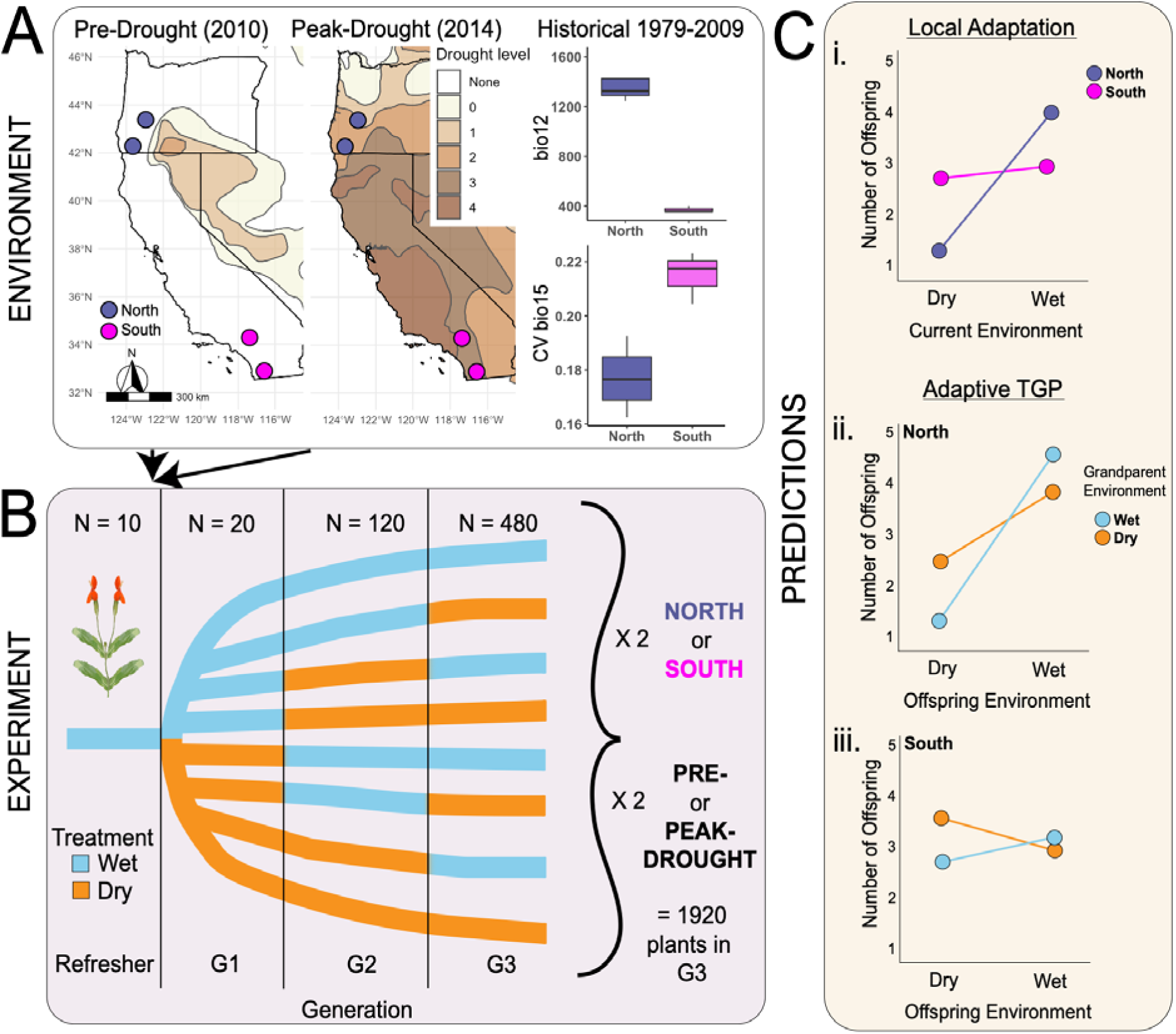
Schematic showing experimental design and predicted relationships. A) Drought severity map for Oregon and California relative to four focal populations of *Mimulus cardinalis* (two northern–blue and two southern–pink) prior to drought (July 13, 2010) and at the peak of the drought (July 15, 2014), as well as boxplots (median, interquartile range, and 1.5 IQR whiskers) of regional historical averages (30-years prior) for annual precipitation (bio12) and interannual variability in precipitation seasonality (coefficient of variation of bio15) for northern and southern populations. Drought levels are not dry (none), abnormally dry (0), moderate drought (1), severe drought (2), extreme drought (3), and exceptional drought (4) and were obtained from the National Drought Mitigation Center, USDA & NOAA.^47^ B) Ten individuals each from northern and southern populations for both pre-drought and peak-drought were refreshed in a greenhouse under wet conditions (which we assumed to be neutral). These plants were then grown for three generations (G1, G2, and G3) in a multifactorial design of wet (blue) or dry (orange) treatments. The end result is eight distinct environmental treatment trajectories (n=1920). C) i) Populations whose native environment reflects the current treatment perform best compared to populations from different climates. ii & iii) adaptive transgenerational plasticity (TGP) is expected to provide an advantage when offspring are exposed to the same environment as their grandparent, but within the context of their local adaptation. Thus, when grandparent and current treatment are mismatched, there is no change in fitness (reflects local adaptation in i).

By measuring phenology, biomass allocation, and components of reproduction, we ask: 1) Does TGP exist?; 2) Does TGP differ across regions?; 3) Is TGP advantageous, resulting in increased offspring fitness when exposed to the same environment?; 4) Does TGP evolve rapidly during severe drought?; and 5) What is the relative contribution of TGP to phenotypic variation compared to other sources, specifically rapidly evolved genetic differences, parental-derived plasticity, and within-generation plasticity? We hypothesized that northern populations that reside in historically less variable climates than southern populations (Fig. 1A) would be more likely to exhibit TGP or at least have more traits that exhibited TGP. We also predicted that TGP would be advantageous if present across both regions, which would express as a reduced difference in local adaptation (Fig. 1C).

## Results

### Existence of TGP and Regional Differences

Grandparent environment significantly affected offspring growth (PC1; p = 0.01; Fig. 2A, Table S1), germination time (p < 0.01; Fig. S1A; Table S1), and flowering date (p = 0.003; Fig. S1B; Table S1), consistent with the influence of TGP on phenotypic traits (Question 1). Droughted grandparents resulted in larger offspring with more flowers and delayed phenology than wet grandparents. Regions differed (Question 2) in the effect of grandparent environment on offspring SLA (Region*Grandparent Treatment p = 0.03; Fig. 2B; Table S1). Offspring of dry grandparents had broader and thinner leaves in the north compared to offspring of wet grandparents, but narrower and thicker leaves in the south. There were also main effects of region, parent treatment, and current treatment (Fig. S1). Southern plants were larger (region p < 0.001; Fig. S1C) and germinated earlier (region p = 0.01; Fig. S1D) compared to northern plants. Offspring in dry treatments were smaller (treatment p < 0.001; Fig S1E), flowered earlier (treatment p < 0.001; Fig S1F), and had lower SLA (treatment p < 0.001; Fig S1G). There were regional differences in current treatment response, resulting in greater plasticity in southern plants for growth (Region*Treatment p < 0.001) and flowering date (Region*Treatment p = 0.02; Fig. S2A,B). Offspring of wet parents had later phenology (germination parent p <0.001; Fig. S1H; flowering date parent p = 0.02; Fig. S1I).

**Figure 2.**
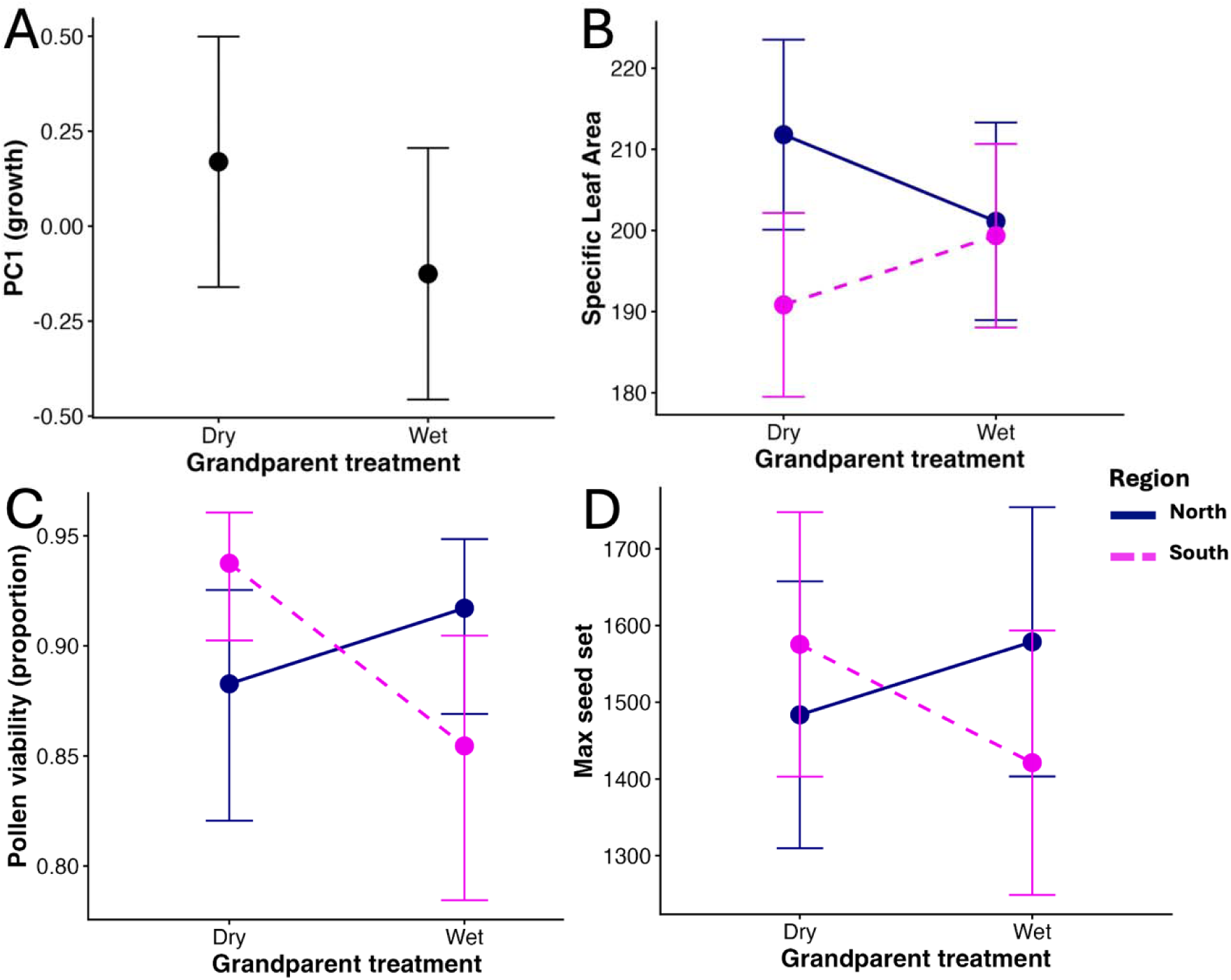
Effect of grandparental environment on offspring growth and fitness. A) PC1 (growth). B) Grandparental environment affects specific leaf area differently between northern (navy solid) and southern (pink dashed) populations. C) Pollen viability is differentially affected by grandparental exposure to wet or dry treatments between northern and southern populations. D) Seed set per flower changes due to grandparental exposure to wet or dry treatments and this differs between northern and southern populations. Points are estimated marginal means from mixed models and error bars are 95% confidence intervals, which includes family-level variation, while treatments are compared within families. Thus significance is not judged by CI interval overlap.

### Adaptive Transgenerational Plasticity

We tested whether TGP was adaptive (Question 3) by examining the fitness components: pollen viability and maximum seed set. Across both models, we found no evidence that grandparental treatment interacted with current treatment (p > 0.05; Table S2), indicating offspring whose grandparents experienced drought did not have an advantage over offspring experiencing novel drought stress, counter to expectations (Fig. 1Cii, iii).

However, we did find strong evidence of regional differences in the way that grandparental treatments affected offspring fitness. Offspring had greater male and female fitness (both Region*Grandparent Treatment p < 0.001) when their grandparents grew in environments that resembled their historical home climates (Fig. 3A,B; Table S2), producing a pattern similar to that of local adaptation (Fig. 1Ci). Offspring from southern populations (dry historical environments; Fig. 1A) had greater reproductive fitness when their grandparents were in dry treatments compared to wet treatments, producing 9.7% more viable pollen (p < 0.001; Fig. S3B) and 10.9% more seeds (p < 0.001; Fig. S3D). Offspring from northern populations (where historical environments are wet; Fig. 1B) had the opposite trend, producing 3.75% less viable pollen (p = 0.07; Fig. 3A) and 6% fewer seeds (p = 0.06; Fig. S3C) when their grandparents experienced dry compared to wet treatments. We also found an effect of treatment (p = 0.002; Fig S4; Table S2), resulting in overall greater reproductive output of offspring in the dry compared to the wet treatment. There was no significant effect of parental treatment on fitness (p > 0.4; Table S2).

**Figure 3.**
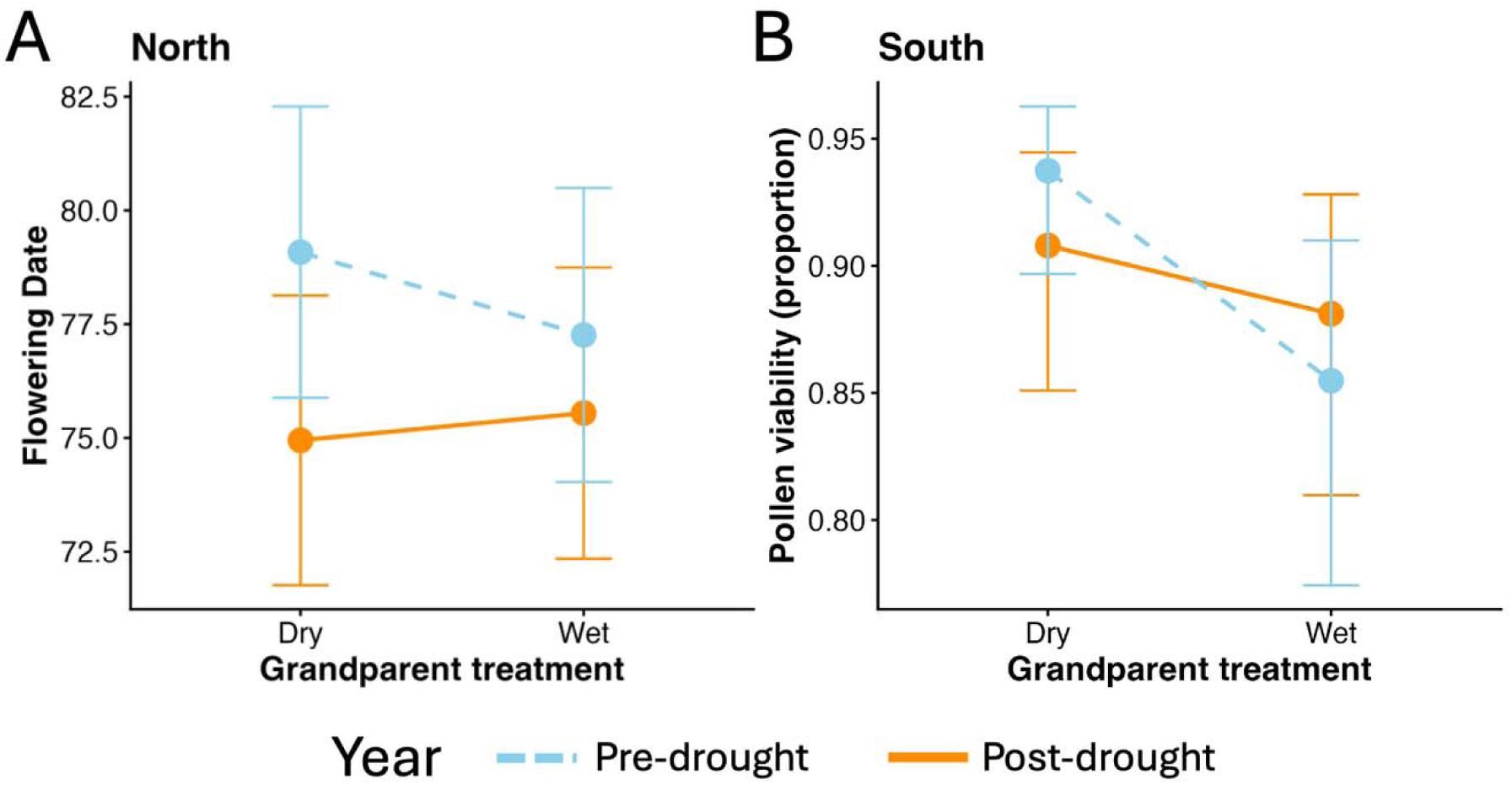
Evolutionary (Pre- and Peak-drought) loss in transgenerational plasticity. A) Effect of grandparent treatment on flowering date in northern plants differs pre-drought versus post-drought. B) Grandparent treatment affects pollen viability in the southern plants pre-drought but not post-drought. Points are estimated marginal means from mixed models and error bars are 95% confidence intervals, which includes family-level variation, while treatments are compared within families. Thus significance is not judged by CI interval overlap.

### Rapid evolution of TGP

To determine whether TGP evolved rapidly during drought (Question 4), we examined if TGP changed between ancestor and descendant populations by testing for significant interactions with Grandparent Treatment and Year (pre-drought and peak-drought) for each region separately. Prior to the drought, flowering date in the north was affected by grandparental treatment (p = 0.03), resulting in later flowering (2 days) when grandparents were in dry treatments. Following the drought, TGP was lost (p = 0.43; Year*Grandparent Treatment p = 0.03; Fig. 3A; Table S3). Similarly, prior to the drought in the south, pollen viability significantly increased when grandparents experienced dry conditions (as stated above; p < 0.001). Following four years of drought, this plasticity was lost (p = 0.12; Year*Grandparent Treatment p = 0.01; Fig. 3B; Table S4). No other traits were found to have significant evolution of TGP.

### Relative effect strength on offspring phenotypes

Lastly, to determine the strength of the effect of TGP compared to genetic change, parental effects, and within-generation plasticity (Question 5), we examined the effect sizes of each within additive mixed models across all nine traits for each region separately (Fig. 4; Table S5). Current treatment generally has the strongest effect on offspring phenotypes (4/9 traits in the north and 5/9 traits in the south having large effect size; es > 0.10). However, grandparental treatment has significantly larger effect than parental treatment (five traits in north; seven traits in south), except for north phenology (germination time and flowering date), and across all traits in the south, parental treatment had no effect (es < 0.01). Grandparental treatment had the greatest effect on branch number (es = 0.03) and pollen viability (es = 0.02) in the north and pollen viability (es = 0.07) and seed set (es = 0.03) in the south. Year had an overall larger effect in the south (four traits) than it did in the north (two traits), corroborating results from our previous experiments.^14,15,16^

**Figure 4.**
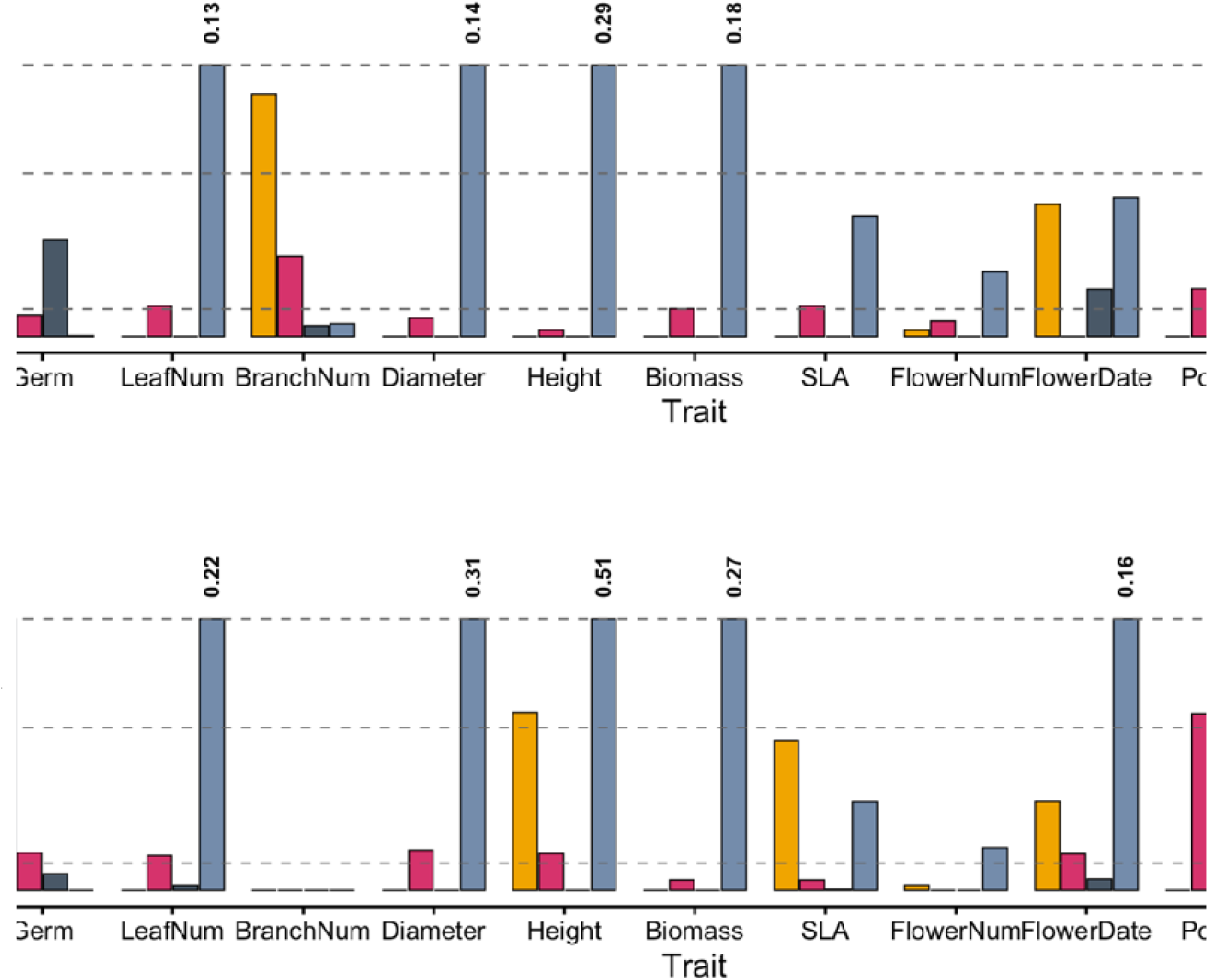
Relative effect of allelic differences (Year), transgenerational plasticity (Grandparent treatment), parental effects, and within-generational plasticity (Current treatment) on plant traits for northern (A) and southern (B) populations. Standardized partial omega-squared effect sizes for germination date (Germ), leaf number (LeafNum), branch number (BranchNum), stem diameter (Diameter), plant height (Height), specific leaf area (SLA), flower number (FlowerNum), flowering date (FlowerDate), pollen viability (Pollen), and maximum seed set (Seed set) as response variables where derived from linear (general linear for pollen) mixed models that consisted of year, grandparental treatment, parental treatment, and current treatment as fixed effects. Germination date does not include within-generation plasticity because offspring had not undergone a treatment yet. Colours indicate year (yellow), grandparent treatment (pink), parent treatment (dark blue), and current treatment (light blue). Dotted lines indicate no effect < 0.01, small effect > 0.01, medium effect > 0.06, and large effect > 0.10. Bars that exceeded 0.10 were capped and their effect size written above.

## Discussion

We found that transgenerational plasticity (TGP), attributed to inherited effects from the environment of the grandparents, significantly affects both male and female fitness, can have greater fitness consequences than allelic or other plastic sources of variation, and can evolve rapidly. These results are the first that we know of to empirically examine TGP in wild plant populations, how it contributes to phenotypic change during rapid evolution to a severe climatic event, and whether it evolves. Our results suggest that the TGP observed in our study likely has an epigenetic origin, because there is no evidence of cumulative effects (no parental or current treatment interactions) and there is rapid evolutionary loss, showing that evolution rapidly acts upon this plasticity. Furthermore, our results suggest two critical errors in the design of plant evolutionary studies, whereby grandparental effects are significantly greater than parental effects across numerous growth and reproductive traits, and that populations respond differently to “benign” refresher generations, showing current practices for removing parental effects are lacking.

We first addressed the existence of TGP and whether its expression differed between regions with different historical precipitation regimes. Theory and empirical studies show that TGP evolves and is adaptive in predictable environments,^2,3,17^ where predictability is inferred by temporal autocorrelation. For example, populations of *Drosophila mojavensis* that occur in predictably variable climates showed greater evolution of TGP than populations with less autocorrelation.^18^ Other studies suggest that TGP evolves where it can prime offspring to perform better in the same stress.^19^ For example, droughted offspring of droughted grandparents or parents had greater growth rates than droughted offspring with no generational history of drought.^20^ In contrast to our prediction that northern populations would exhibit TGP while southern populations would not (based on climatic variability; Fig. 1A), we found TGP in both regions and stronger TGP in the south. It is possible that interannual variability in precipitation seasonality is an inadequate metric of climatic predictability, and perhaps the southern region would be considered more stable with regard to other, more biologically relevant climate, variables. However, southern populations have higher rates of within-generation plasticity in phenotypic traits,^15^ which is generally attributed to heterogeneous environments (i.e., high rates of variability). Diaz et al. (2021) found that more variable *Drosophila mojavensis* climate was also more predictable and this was associated not only with TGP evolution but also with within-generation plasticity.^18^

While we did not find regional differences in TGP that matched our predictions based on precipitation variability, we did find that TGP is both adaptive and maladaptive (Question 3), the direction of which is environmentally dependent across regions. One way that stress can be passed on across multiple generations is in the form of strong cumulative effects, through multiple generations of stress occurrence.^20^ For example, three generations of harsh temperatures resulted in significantly lower fitness in *Arabidopsis* and different phenotypic outcomes occurred depending on the number of generations exposed.^4,21^ Similar to mutation accumulation, recurrent stress across generations can seemingly result in an accumulation of maladapted epialleles. This is not always the case, because cumulative effects can also result in greater priming, such as when three generations of drought resulted in greatest seedling development in the herb *Polygonum persicaria*.^20^ Furthermore, Groot *et al.* (2016) conducted a study on salt stress across three generations in an inbred line of *Arabidopsis* and found that salt stress in the parental generation was maladaptive, but was alleviated when both grandparents and parents experienced salt stress.^22^ Surprisingly, we find no interactions between grandparent treatments and current or parent treatments (Fig. 2). Instead, we observe the lasting effects of grandparent environment on offspring phenotypes even with an intervening parental generational in different conditions.

Indeed, here we find that the fitness direction was determined by the grandparent treatment regardless of the current or parental treatment entirely. Similar to local adaptation with fitness tradeoffs, we find that TGP is advantageous *only* when the environment of the grandparent matches historical climate, regardless of the environment of the offspring (Fig 2B-D). This is contrary to how TGP is generally expected to work (above; Fig. 1Cii,iii).^6^ Stress-induced changes in resource provisioning during embryo development is an alternative to cumulative effects that could result in long-lasting fitness consequences. In this scenario, droughted grandparents allocate more stress hormones and less resources to developing seeds, resulting in the next generation acquire fewer resources, leading to the same reduced provisioning for grand- offspring.^23^ In contrast, the Silver Spoon hypothesis states the opposite, where greater provisioning to developing embryos in favourable environments can result in greater success across multiple generations.^23^ Our study reflects this outcome; grand-offspring had greater reproductive success when grandparents experienced a favourable environment, while they paid a fitness penalty when grandparents had experienced a mismatched environment. Further research examining fitness consequences across each generation is necessary to better understanding this potential mode of TGP.

Importantly, however, we find evidence that TGP can be rapidly labile to stress perturbation, resulting in an evolutionary loss of TGP in two traits following severe drought (Fig. 3). Prior to an historic drought, populations in the north exhibited a two-day difference in flowering date depending on whether grandparents experienced dry versus wet environments, while post-drought descendants no longer exhibited TGP. Similarly, pre-drought ancestors in the south had ∼10% more viable pollen when grandparents were in dry compared to wet environments, which was lost in peak-drought descendant populations. Thus, not only do we have evidence from our prior studies of rapid allelic turnover contributing to evolution in drought responses,^14,16^ we also find rapid evolution of TGP in the wild. These results provide evidence that there could be selection against TGP as climatic variability increases. Furthermore, the evolutionary loss of TGP provides evidence that the pattern we observed could be epigenetic in origin rather than Silver Spoon. While we cannot completely rule out lagging Silver Spoon effects, this type of effect is expected to be stable across time, thus would not be rapidly labile compared to epialleles that are.^7^ We are not aware of any other studies that has shown that TGP has the capacity to rapidly evolve in the wild.

Teasing apart the main source of phenotypic variation (i.e., G x E_TGP_ x E_parent_ x E_Within-generation plasticity_) has been of great debate since the recognition of TGP.^5,6^ While parental effects are ubiquitous within ecology and evolutionary studies, our results suggest more emphasis should be placed on grandparental or perhaps great-grandparental environments than parental environments. Indeed, we find that grandparent environment has much stronger effect than parental effects in almost every trait examined. Furthermore, the significant effect that TGP has on pollen viability and seed set in our study, particularly for southern populations, has the potential for great evolutionary consequences. Highly conserved epigenomic modifications have been associated with pollen development and seed production in *Arabidopsis*.^24,25^ Furthermore, epiallelic variation diversifies at greater rates than genetic mutation,^7^ and therefore contributes a significant role for phenotypic differences and potentially allows for faster evolution.^1,26–29^ Although we do not know if the source of TGP in our study is epigenetic, our results show that TGP had a greater effect on fitness components than recently evolved allelic differences in southern populations (Fig. 4) and a similar effect for pollen viability in northern populations. Thus the differences in the rate of rapid evolution (faster in the south and slower in the north) that we found previously,^14,15,16^ could be a consequence of differences in fitness associated with TGP between these regions during the drought. Our results are the first that we know of that describe grandparentally-derived plasticity responding in this manner.

Our results provide new insight on how phenotypic variation arises through transgenerational plasticity. Grandparental environments can have significant consequences for both male and female fitness, and this work emphasizes the need to incorporate direct fitness effects into studies on transgenerational plasticity and epigenetics. While we do not know the mechanistic source of the TGP presented here, that it is labile across time and does not interact with the parental or current environments, suggests it is likely epigenetic in origin. Future work should probe mechanistic sources of this result. We find that the capacity to express transgenerational plasticity and the significance of that expression is genetically determined (regional differences) and that it is only adaptive when conditions reflect historical climate patterns, but is maladaptive when conditions do not match. Importantly, our results show that grandparental effects can be much stronger than parental effects and thus have significant implications for how we design experiments to reduce environmental effects. We show clear evidence that future studies on transgenerational plasticity should examine the phenotypic consequences of environments from generations beyond the parental environment and that we need to re-evaluate how refresher generations are conducted to obtain accurate genetic phenotypic differences.

## Materials and Methods

### Study system

*Mimulus cardinalis* Spach *(Erythranthe cardinalis* Doughl. ex Benth)^30–34^ is an herbaceous, hummingbird-pollinated perennial plant that occurs within riparian areas across southwestern North America from central Oregon to northern Baja California.^34^ Populations (Table S6) in the northern portion of the range experience higher mean annual precipitation and relatively lower precipitation variability across years compared to populations in the south, which experience lower mean annual precipitation and relatively more variable precipitation patterns across years (Fig. 1A).^14^ From 2012-2015, a record-breaking 4-year drought impacted the entire range.^35^ Seeds of *M. cardinalis* were collected prior to the drought onset in 2010 (ancestral genotypes) and collected at the peak of the drought (either 2014 or 2015, depending on the drought peak for that site), representing post-drought descendant genotypes.^14,15^ Most individuals of this species reproduce for only one-two years before dying,^8^ therefore it is unlikely that seeds collected from the descendant generation are from the same plant that ancestral seeds were collected from.

### Growth Conditions

Wild seeds were refreshed in a common greenhouse with 27/15°C day/night temperatures, 12-hour days, and saturating water.^14^ Germination of these seeds was not affected by the invisible fraction problem.^14,36^ We used the resurrection approach^13^ to compare ancestor and descendant progeny.

Refreshed seeds were sown in 233ml square pots filled with a 2:1 mix of fine-grade sand (QUIKRETE® Premium Play Sand® No. 1113) and Sunshine Mix #4 without arbuscular mycorrhizal fungi (AMF; Sun Gro Horticulture, Agawam, MA), with 0.52g per pot of evenly distributed Osmocote Plus slow-release fertilizer (Scotts Co LLC, Marysville, Ohio). Seeds were misted for two minutes, four times a day until germination. Afterward plants were bottom watered to field capacity until established, characterized as the development of three sets of leaves. Plants were then either placed in a wet treatment (maintained at field capacity for the experiment duration) or a dry treatment (held just above wilting point). To ensure all plants experienced the same level of abiotic stress, the dry treatment was tailored to each plant’s wilting point, which was visually assessed multiple times per day; water was supplemented when wilt occurred. Greenhouse conditions were set to 14-hour day, 28/18°C day/night temperatures, with natural light being supplemented by LED lights as needed. Plants were grown in their respective treatments through fruit maturation; thus, the duration of the experiment differed depending on individual plant ontogeny. This ensured that full seed development occurred within the treatment conditions. The same growth conditions and treatment categories were used across all three generations.

### Establishment of generations

Five full-sib families (seeds derived from a single fruit) from each refreshed population and year were selected randomly and then introduced to the two treatments (grandparental; n = 80; Fig.1B). First flowers were self-pollinated during their treatment and fruits matured within the treatment. To replicate full-siblings, six seeds from each grandparental generation plant were sown and then placed in respective treatments (three in wet and three in dry) to establish the second generation (parental; n = 480; Fig.1B). First flowers were again self-pollinated and fruits matured under treatment. For the final generation, four seeds from each parent plant were sown and then two each were grown in wet or dry treatment (offspring; n = 1920; Fig. 1B).

### Measurements

In each generation, pots were checked daily and the date of germination/emergence was recorded. At the time of first flower, the most recently fully expanded leaf not subtending the flower was removed and measured for specific leaf area (cm^2^/g; SLA). SLA was calculated as the surface area of one side of the leaf divided by the dry mass of the leaf. We measured projected area of freshly harvested leaves with a LI-300C (LiCor Inc., Lincoln, NB) and then weighed with XPR Micro balance (Mettler-Toledo AG, Greifensee, Switzerland) following drying in a 50°C oven for two days. We recorded date of first flower opening and, after fruit ripening, plant height, stem diameter at base, number of branches, and number of leaves. Plants that had not flowered after 120 days (∼4 months) were not included in flowering date. This cut off was determined based on typical growing season length. Pollen was collected from half of the individuals in the third generation from one undehisced mature anther from the first flower and placed in Alexander’s Triple Stain.^37^ For an approximately 300 pollen grains per individual, we counted the number aborted versus mature grains, we used the number of mature grains compared to total grains as a proxy for percent viable. The first flower was saturated with pollen by placing a dehiscing anther fully on the stigma and pressing the stigma together to dislodge pollen. Stigmas were monitored the following day to ensure successful fertilization had been complete, inferred when stigmas stayed closed instead of reopening. At maturation, we collected fruits and weighed total seed set. We determined an average seed weight (0.023mg) by counting and weighing the total seeds produced in 444 fruits for individuals across all three generations. Seed totals were then estimated based on their total weight relative to the average seed weight.

### Statistical Analyses Data curation

Here we analyzed data derived from the offspring generation (after three generations of treatments). To distill multiple traits into fewer axes of trait variation, we used principal components analysis to ordinate multivariate data. We first examined pairwise correlations (*cor* function; Fig. S5A)^38^ between all pairs of the seven leaf and phenological traits (excluding fitness component traits: pollen viability and maximum seed set) to examine possible trait redundancy (r > 0.85; no variables were excluded) and coherence (Kaiser-Meyer-Olkin MSA < 0.6), which tests whether samples are adequately associated to one another to be used in a PCA.^39^ Flowering date (MSA = 0.32) and specific leaf area (0.43) were removed, leaving five traits for the PCA. Some data required transformation (log of germination and square-root biomass) prior to conducting a PCA (Fig. S5B) and all data was scaled within the *prcomp* function.

### Treatment interactions and model set up

Our full design had the potential for 4- and 5-way interactions that would be difficult to interpret and were not of direct relevance for answering our questions. So, we did the following to ensure that the factor of least relevance (parental environment) could be included exclusively additively without distorting or obscuring important patterns. We first filtered the data by year (pre-drought ancestors) and split by region to conduct linear mixed effects models using the *lmer* function from ‘lme4’ package^40^ with 3-way treatment interactions (grandparent, parent, and current; Table S7). Since each additional model introduced below, has a different model structure, we also confirmed that using an additive parental treatment structure instead of an interacting one did not distort grandparental estimates and effects by comparing the four-way model with the additive parental structure using likelihood ratio test (LRT; Table S8).

### Existence of TGP and Regional Differences

We examined all response variables, except for fitness responses, to address whether TGP exists (Question 1) and if there are differences in TGP between regions (Question 2). We examined baseline ancestral samples only (i.e., dataset filtered to year=pre-drought; 594 observations) and included region, grandparental treatment, current treatment and their interactions as fixed effects. Replicate and parental treatment were non-interacting fixed effects, while grandparental ID and site nested within region were random effects. Prior to building each model, orthogonal contrasts for each predictor were ensured by setting the *contrasts* to ‘*contr.sum’*. We examined each model for homoscedasticity and normality using qqplot (generated from *qqmath* function in the ‘lattice’ package)^41^ and histograms of the residuals (*hist* function in base R). To test model significance, we assessed Type III ANOVA with Kenward-Roger approximated degrees of freedom (lmerTest). When interaction terms were significant (p < 0.05), we conducted planned pairwise comparisons using *emmeans, pairs,* and *contrast*.^42^ Figures depicting estimated marginal means were generated in ‘ggplot2’.^43^

### Adaptive Transgenerational Plasticity

To address whether TGP is adaptive (Question 3) we performed the same analyses as above but for the two fitness component response variables: pollen viability (male fitness; 331 observations) and seed set (female fitness; 820 observations). Because pollen viability was zero-inflated with differing total pollen grains per sample, we used a *glmer* with a binomial distribution and a Wald chi-square Type-III *Anova* (‘car’ package) for significance testing.^44^ To test for sample dispersion and model fit, we used *testDispersion* from the ‘DHARMa’ package,^45^ which required adding observation-level random effect (OLRE). Maximum seed number followed a normal distribution and thus *lmer* was used. Generally, for TGP to be considered adaptive, the grandparental environment would increase the fitness and response of offspring to the same stimulus, thus an interaction between TGP*Treatment would be expected (Fig 1Cii, iii).

### Rapid evolution of TGP

To address whether TGP evolves (Question 4) we included data derived from both time points (pre-drought ancestrors and peak-drought descendants) and split the data by region (South n = 807; North n = 761). We then used the same model structure, including Rep and parental treatment as main effects, to examine differences in all response variables (PC1, germination time, flowering time, SLA, pollen viability, and seed set). Here interacting terms were year (pre- and peak- drought), grandparent treatment, and current treatment.

### Relative effect strength on offspring phenotypes

Lastly, we used regionally subdivided datasets to assess the relative effect size (Question 5) of TGP, allelic (yearly) differences, parental-derived plasticity, and within-generation plasticity. Here we examined the main effects (partial omega-squared from the ‘effectsize’ package)^46^ from additive mixed effects models on all nine (including traits in the PCA; leaf number, stem diameter, branch number, flower number, flowering time, SLA, germination time, pollen viability, and seed set) traits.

## Data and Code availability

Data from this manuscript can be found at X and code for analyses found at Y. [To be uploaded to Dryad and GitHub once accepted].

## Author Contributions

HAB designed the experiment with ALA. HAB conducted the experiment and collected the data with help from DRM. HAB analyzed the data and wrote the manuscript with contribution from ALA and edits from DRM.

## Acknowledgments

We thank Dr. Katie Marshall for feedback on the initial experimental design. Many individuals assisted in data collection: Simi Badyal, Cara-Lynn Branch, Sam Dang, Lydia Fong, Julien Grebert, Carly Hilbert, Amy Liu, Hayat Mahdjoub, Annie McLeod, Stephanie McPhail, Bryn Murphy, Burak Ozkan, Gundeep Pannu, David Qi, Maya U Schueller Elmers, Mia Waters, Maggie Yang, and Noah Zovickian. We thank Victoria Sork for feedback on previous iterations of the manuscript. Funding for this research was provided by National Science Foundation (NSF) PRFB 2305993, Natural Sciences and Engineering Research Council (NSERC) of Canada PDF 578168, NSERC PGS-D, and Rick Hansen Man in Motion Fellowship from the University of British Columbia to H.B., and NSERC Discovery grant AWD-010335 to A.A.

